# Open-source mapping and variant calling for large-scale NGS data from original base-quality scores

**DOI:** 10.1101/2020.12.15.356360

**Authors:** Olga Krasheninina, Yih-Chii Hwang, Xiaodong Bai, Aleksandra Zalcman, Evan Maxwell, Jeffrey G. Reid, William J. Salerno

## Abstract

Standardized genome informatics protocols minimize reprocessing costs and facilitate harmonization across studies if implemented in a transparent, accessible and reproducible manner. Here we define the OQFE protocol, a lossless read-mapping protocol that retains key features of existing NGS standard methods. We demonstrate that variants can be called directly from NovaSeq OQFE data without the need for base quality score recalibration and describe a large-scale variant calling protocol for OQFE data. The OQFE protocol is open-source and a containerized implementation is provided.

## Introduction

Public genomic initiatives such as the UK Biobank^1^, 1000 Genomes Project^2^, and the Human Diversity Project^3^ have established study populations and data resources that support a wide range of research, from human health to technology development, both as standalone data and as large-scale assets to complementary research programs. Given the size of these datasets, increasingly measured in hundreds of thousands of samples and petabytes of data, routine or custom reprocessing is infeasible for all but the most well-resourced users. However, such reprocessing is inevitable in the long term as methods improve and the data sets themselves grow.

Standardized genomic data protocols can obviate reprocessing by allowing users to harmonize their own data with large resources, ensuring the interoperability of datasets. In 2017, researchers defined functionally equivalent (FE) pipelines for sequence read mapping that were implemented across multiple large-scale sequencing projects, harmonizing more than 400,000 whole-genome samples worth of data with a three-fold reduction in size, achieved largely through lossless reference-based compression (CRAM) and a lossy quality-score binning from the native HiSeq X 8-value scheme to a recalibrated 4-value scheme^4^.

The original quality functionally equivalent (OQFE) protocol presented here adapts the FE protocol so that the original raw read data (i.e. FASTQ files) can be recovered from the resulting CRAM files. Applied to NovaSeq data, which have natively 4-valued quality scores, OQFE CRAMs are comparably sized. Minor updates of constituent programs are made to resolve known issues. Variants can be directly called from these CRAM files, as demonstrated with the DeepVariant^5^ and GLnexus^6,7^ protocol described below.

## Methods

### OQFE Protocol

The OQFE protocol maps raw reads (FASTQ) with BWA-MEM to the GRCh38 reference in a deterministic manner, retaining all supplementary alignments. Mate tags are added with samblaster as specified in the FE protocol. OQFE CRAMs contain all reads from the input FASTQs and meet all FE tag specifications. Duplicate reads are then marked with Picard 2.21.2, which resolves a known issue with the FE version of Picard (2.4.1), in which the representative read in a set of duplicate reads can depend on the sequence input order, potentially resulting in an order-dependent set of supplementary alignment duplicates. The final OQFE CRAM is compressed with samtools, without any base quality score recalibration or binning. OQFE CRAMs are thus forward compatible with the FE quality score recalibration and binning steps. Table 1 details the software versions, references and commands for each step and notes differences from the FE protocol.

**Table 1.**

| FE Requirements | OQFE protocol https://hub.docker.com/r/dnanexus/oqfe |  |  |  |
| --- | --- | --- | --- | --- |
|  | Program | Version | Command Options | OQFE Update Notes |
| align reads: GRCh38DH with .alt file, BWA mem v0.7.15 -Y -K 100000000 | bwa<sup>14</sup> mem | 0.7.17 | -K 100000000 -Y |  |
| Retain the minimal set of tags (RG, MQ, MC and SA). NOTE: an additional tool may be needed to add the MQ and MC tags if none of the tools add these tags otherwise. One option is to pipe the alignment through samblaster with the options -a --addMateTags. | samblaster<sup>15</sup> | 0.1.24 | --addMateTags -a | Picard FixMateInformation adds mate tags as required and adjusts other mate information (e.g. mate chromosome and position, insert size, bitflag) in ways not specified or required by FE protocol. |
| Accurate duplicate marking of supplementary alignments by Picard requires mapped reads to be name sorted. | sambamba<sup>16</sup> sort | 0.6.4 | -n |  |
|  | sambamba merge | 0.6.4 |  |  |
| mark duplicates: Picard v2.4.1 or above | picard<sup>17</sup> MarkDuplicates | 2.21.2 | ASSUME_SORT_ORDER=queryname | Resolves a known issue<sup>18</sup> concerning which reads in a duplicate set are marked as a duplicate, which can affect the number of supplementary duplicates. |
| Coordinate-sorted CRAM | sambamba sort | 0.6.4 |  |  |
| BQSR |  |  |  | Excluded in OQFE |
| apply BQSR: 4-bin |  |  |  | Excluded in OQFE |
| convert to CRAM: PG records; RG: PL, PU, SM, LB; tags: RG, MQ, MC, SA, original query names | samtools<sup>19</sup> view | 1.9 | -C |  |

### OQFE DeepVariant Protocol

Variants were called on each CRAM with DeepVariant^5^ 0.10.0 using a deep learning model retrained on exome data sequenced with the same protocol as was used to sequence the UK Biobank samples^8^. Variant calls were restricted to the exome capture region and the 100 base-pairs flanking each capture target, resulting in a gVCF (genomic VCF) for each sample containing all variant genotypes and compressed representations of reference regions without called variant genotypes.

The OQFE protocol was applied to the 200,000 UK Biobank (UKB 200K) exome samples^9^ with the containerized OQFE pipeline (https://hub.docker.com/r/dnanexus/oqfe). Per-sample gVCFs were generated via the DeepVariant protocol described above and merged with GLnexus 1.2.6 using the default ‘DeepVariantWES’ parameters^6,7^. Table 2 provides exact commands and access to all required resource files.

**Table 2.**

| Program | Version | Command Options |
| --- | --- | --- |
| DeepVariant<br>*For UKB 200K, DeepVariant was executed with a GPU-accelerated implementation that produces identical gVCF calls relative to the open-source version | 0.10.0 | --ref [Reference]<br>--regions [BED file]<br>--customized_model [model file]<br>--output_gvcf |
| GLnexus | 1.2.6 | -i config=DeepVariantWES<br>-i bed_ranges_to_genotype [BED file] |
| Reference: ftp://ftp.ncbi.nlm.nih.gov/1000genomes/ftp/technical/reference/GRCh38_reference_genome/<br>BED file: xgen_plus_spikein_100bpBuff.GRCh38.bed (See Supplemental Data)<br>Model: model.ckpt-22236.tar (See Supplemental Data) |  |  |

### HG002 benchmark data

Two sets of NovaSeq exome sequence data were generated from the HG002 control sample^10^ via the exome sequencing protocol applied to UK Biobank samples^8^ and then mapped via the OQFE protocol. Two additional CRAMs were created from each HG002 OQFE CRAM by recalibrating the base qualities (+BQSR CRAM) and then applying the FE binning strategy (+BQSR+FEbin CRAM) as described in the FE protocol^4^. Original quality scores are not retained in either type of derived CRAM. All HG002 CRAMs were called with the OQFE DeepVariant protocol within the exome capture regions and evaluated with hap.py 0.3.8^11^ against the Genome In A Bottle HG002 high-confidence variants (v3.3.2) within the corresponding HG002 high-confidence regions^12,13^.

## Results

NovaSeq OQFE CRAMs retain original quality scores with only a modest increase in size (10-12%) compared to FE CRAMs and are approximately one-third the size of CRAMs with recalibrated quality scores (Table S2). The UKB 200K exome CRAMs (n=200,643) average 838 MB per sample, totaling approximately 174 TB. Compared to native NovaSeq data (i.e. read-name-sorted and compressed FASTQs), OQFE CRAMs maintain the three-fold reduction in size offered by FE CRAMs (Table S2).

To demonstrate that variants can be called directly from NovaSeq exome OQFE CRAMs without a loss of quality, we compared HG002 variant performance at two coverages (45x and 70x) and three quality score binning strategies: OQFE (native 4-valued), +BQSR (40-valued), +BQSR+FEbin (non-linear 4-valued). On average 21.6k SNVs and 880 indels per CRAM were compared to 22,587 high-confidence HG002 variants (21,675 SNVs and 912 indels), providing precision, recall and F1 scores for each of the six experiments (Table S1). As shown in Figure 1, variant performance varies more with coverage and with variant type than it does with quality-score binning. Summing across variant types and coverages (Table S1), we observe that OQFE has slightly fewer false negatives (FN=384) and false positives (FP=127) than each +BQSR (FN=390, FP=130) and +BQSR+FEbin (FN=394, FN=163).

**Figure 1:**
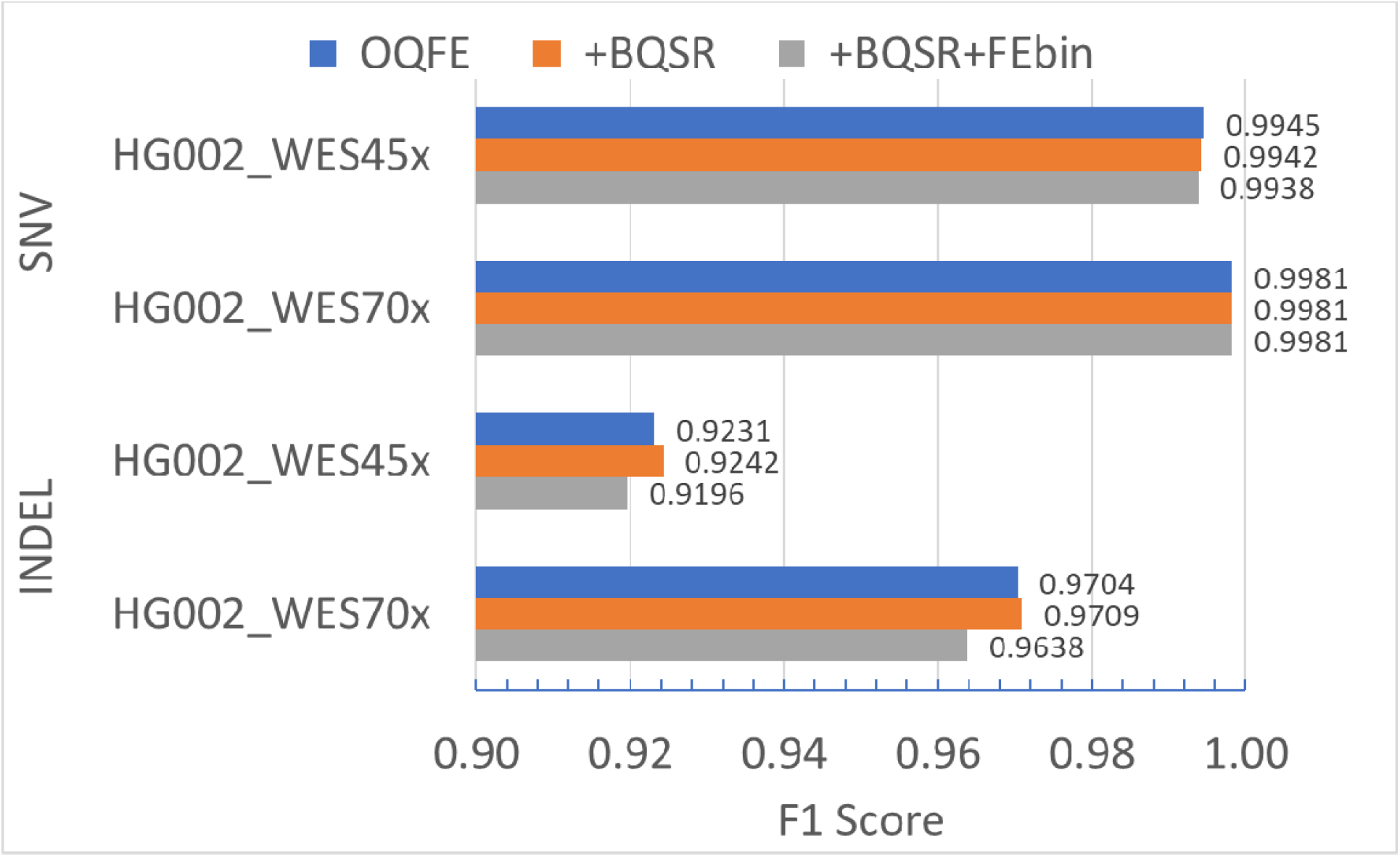
Comparison of quality score binning strategies by variant type. F1 scores for SNVs differ by less than 0.0008 between all binning strategies at each coverage, and OQFE indel F1 scores are within 0.0012 of +BQSR values.

## Discussion

Public genomic data represent significant investments in money, human effort and subject participation, all of which demand the data be both equitably actionable in the short term and durable in the long term. The size and complexity of public genomic datasets present a barrier to many users, even if the data are freely accessible. At the same time, the data must be amenable to current and future research that requires reprocessing (e.g. a new reference genome). While OQFE CRAMs are lossless relative to FASTQs and thus a durable long-term resource, they are reference-coordinate sorted and compressed. If FASTQs are required, we recommend OQFE CRAMs be name-sorted prior to conversion to avoid reference-specific correlation in the FASTQ read order.

We also note that the OQFE protocol avoids potential overbinning of NovaSeq quality scores. The FE protocol assigns all recalibrated quality scores greater than 23 (PHRED scale) a value of 30. When applied to the native NovaSeq quality scores of 2, 11, 25 and 37, the FE binning would both fail to distinguish between the two highest quality scores and deflate the highest value. While the OQFE+DV results described here are largely similar across quality-score binning strategies, we recommend that users with NovaSeq data evaluate any quality-score processing with respect to their variant calling protocol prior to analysis.

Lastly, we recognize that the cost to egress, store and reprocess data is compounded by the expertise required to maintain, optimize and execute genomic software at scale. To this end, all methods described here rely only on open-source software, and we provide a single containerized OQFE pipeline with all required source and validation files that can be executed on any local or cloud infrastructure that supports Docker containers. This ‘open-source-first’ policy combined with standardized descriptions ensures that users can execute these exact methods autonomously on standard hardware while also enabling commercial providers to facilitate accelerated and at-scale processing with specialized technology.

## Supporting information

Custom DeepVariant Model

WES Capture BED

OQFE Supplementary Tables

